# TACN (1, 4, 7-Triazacyclononane) restores the activity of β-lactam antibiotics against Metallo-β-Lactamase producing *Enterobacteriaceae*: The exploration of potential Metallo-β-Lactamase inhibitors

**DOI:** 10.1101/366146

**Authors:** Anou M. Somboro, Daniel G. Amoako, John Osei Sekyere, Hezekiel M. Kumalo, René Khan, Linda A. Bester, Sabiha Y. Essack

**Affiliations:** Antimicrobial Research Unit, School of Health Sciences, University of KwaZulu-Natal, Durban, South Africa; Biomedical Resource Unit, College of Health Sciences, University of KwaZulu-Natal; Durban, South Africa; Department of Medical Microbiology, Faculty of Health Sciences, University of Pretoria, South Africa; Discipline of Medical Biochemistry, School of Laboratory Medicine and Medical Science, University of KwaZulu-Natal, Durban, South Africa

**Keywords:** TACN, Metallo-β-lactamase inhibitor, binding affinity, β-lactams, *Enterobacteriaceae*

## Abstract

Metallo-β-lactamase producing *Enterobacteriaceae* are of grave clinical concern particularly as there are no Metallo-β-lactamase (MBL) inhibitors approved for clinical use. The discovery and development of MBL inhibitors to restore the efficacy of available β-lactams are thus imperative. We investigated a zinc-chelating moiety, 1, 4, 7-triazacyclononane (TACN) for its inhibitory activity against clinical carbapenem-resistant *Enterobacteriaceae*. Minimum inhibitory concentrations (MICs), minimum bactericidal concentrations (MBCs), serum effect, fractional inhibitory concentrations index and time-kill kinetics were performed using broth microdilution techniques according to the Clinical Laboratory Standard Institute (CSLI) guidelines. Enzyme kinetic parameters and cytotoxicity effects of TACN were determined using spectrophotometric assays. The interactions of the enzyme-TACN complex were investigated by computational studies. Meropenem regained its activity against carbapenemase-producing *Enterobacteriaceae*, with the MIC decreasing to 0.03 mg/L in the presence of TACN. TACN-Meropenem combinations showed bactericidal effects with MIC/MBC ratio of ≤4, and synergistic activity was observed. Human serum effects on the MICs were insignificant, and TACN was found to be non-cytotoxic at concentrations above the MIC values. Computational studies predicted that TACN inhibits MBLs by targeting their catalytic active site pockets. This was supported by its inhibition constant K_i_ = 0.044 µM and inactivation constant k_inact_= 0.0406 (min^-1^) demonstrating that TACN inhibits MBLs efficiently and holds promise as a potential inhibitor.

**Importance:** Carbapenem-resistant *Enterobacteriaceae* (CRE)-mediated infections remain a significant public health concern and have been reported as critical in the World Health Organization’s Priority Pathogens List for the Research and Development of New Antibiotics. CRE produce enzymes such as Metallo-β-lactamases (MBLs), which inactivate β-lactam antibiotics. Combination therapies involving a β-lactam antibiotic and a β-lactamase inhibitor remain a major treatment option for infections caused by β-lactamase-producing organisms. Currently, no MBL inhibitor-β-lactam combination therapy is clinically available for MBL-positive bacterial infections. Hence, developing efficient molecules capable of inhibiting these enzymes could be a promising way to overcome this phenomenon. TACN played a significant role in the inhibitory activity of the tested molecules against CREs by potentiating the carbapenem. This study demonstrated that TACN inhibits MBLs efficiently and holds promises as a potential MBLs inhibitor to help curb the global health threat posed by MBL-producing CREs.

## Introduction

β-lactams are broad-spectrum antibiotics with relatively minimal toxicity, making them antibiotics of choice for many bacterial infections(1). However, the surge in resistance to β-lactams in Gram-negative bacteria is affecting the management of infections worldwide(2). β-lactams are rendered ineffective through different mechanisms, including reduced membrane permeability through porin down-regulation and efflux, target site modification, and most importantly, the production of β-lactamases, enzymes that hydrolyze β-lactams(3).

Metallo-β-lactamases (MBLs) are Ambler class B β-lactamases with zinc ions at their active sites. The zinc ions orient a hydroxide nucleophile to attack the carbonyl on a β-lactam ring(4, 5). Sub-class B1 MBLs are the most clinically relevant among the class B MBLs. They possess a binuclear active site, which contains either one or two Zn (II) ions and hydrolyze all β-lactam antibiotics in clinical use except monobactams(6). MBLs such as New Delhi Metallo-beta-lactamase (NDM), Verona Integron-Borne Metallo-β-Lactamase (VIM) and Imipenemase (IMP) mediate high-level resistance to antibiotics in carbapenem-resistant *Enterobacteriaceae* (CRE)(7). CRE have been identified across the globe, including in all African regions(8–10) and have been ranked as critical in the World Health Organization’s Priority Pathogens List for the Research and Development of New Antibiotics(11). It is therefore imperative to invest in the discovery and development of MBL inhibitors (MBLIs) for combination therapy with carbapenems.

Potential MBLIs investigated to date have shown substantial *in vitro* activity but cannot be used clinically due to the simultaneous inhibition of human metallo-enzymes or their cytotoxicity effects(12, 13). MBLIs such as aspergillomarasmine A (AMA)(13), Ca-ethylenediaminetetraacetic acid (Ca-EDTA)(12), 1,4,7-triazacyclononane-1,4,7-triacetic acid (NOTA) and 1,4,7,10-tetraazacyclododecane-1,4,7,10-tetraacetic acid (DOTA)(14) are the few non-toxic MBLIs identified to date and act by chelating the zinc ions(15). Herein, we investigated the inhibitory activity of 1, 4, 7-triazacyclononane (TACN) against well-characterized sub-class B1 MBL-producing *Enterobacteriaceae*, and to the best of our knowledge, this is the first time TACN has been investigated for its inhibitory activity against MBL-producing *Enterobacteriaceae* to potentiate the activity of meropenem (MEM). TACN has a different chemical structure with the smallest molecular weight in the family of metal chelator series reported. We believe that exploring and understanding the mechanisms of biologically relevant chemical moieties is the surest way to find clinically potent MBL Inhibitors that will give a lasting solution to this global menace.

## Results

### Minimum inhibitory concentrations (MICs), minimal bactericidal concentrations (MBCs), and MIC/MBC ratio

TACN is a cyclic organic compound within the NOTA and DOTA series and derived from cyclononane by replacing three equidistant CH_2_ groups with NH groups(16) (Figure 1). It can be seen from the structures that TACN is unique from that of NOTA and DOTA (family of compounds) (Figure 1). The results revealed that this molecule (TACN) successfully restored the activity of MEM against characterized clinical isolates of carbapenem-resistant *Enterobacteriaceae* expressing acquired sub-class B1 metallo-beta-carbapenemases and reference strains with MIC values as low as 0.03 mg/L shown in table 1 and 2. Also varying inhibitory concentrations of TACN considerably enhanced the efficacy of MEM against the MBL-producing bacteria (varying from 2 mg/L to 64 mg/L). The lowest concentration of TACN at which most of the MEM activity was restored was 8 mg/L (Table 1 and 2). Therefore, all subsequent experiments were conducted with a fixed concentration of 8 mg/L (6.25%) of TACN. None of the tested serine β-lactamases (SBLs) types (*K. pneumonia* OXA-48 and *E. cloacae* KPC-2) was affected by TACN, substantiating the substrate spectrum of this compound (Table 1). MBCs of MEM were determined in the presence of TACN at a fixed concentration, and the MBC/MIC ratio of the combination ranged from 1-to 4-fold against both reference and clinical CRE isolates used in this study, except two clinical isolates, which had 8x the MIC values (Table 2). *K. pneumoniae* ATCC BAA 1706 and *E. coli* ATCC 25922 had no changes in their MIC values when challenged with MEM alone and MEM+TACN.

**Figure 1:** Chemical structures of [1]: 1, 4, 7-triazacyclononane (TACN), [2]: 1,4,7-triazacyclononane-1,4,7-triacetic acid (NOTA) and [3]: 1,4,7,10-tetraazacyclododecane-1,4,7,10-tetraacetic acid (DOTA).

**Table 1:** Inhibitory activity of TACN and MEM alone and in combination against reference strains of carbapenem-resistant Enterobacteriaceae.

| Bacterial species | Metallo-β-lactamase produced (Class B) | MIC (mg/L) |  |  |  |  | MBC (mg/L) | MIC/MBC (mg/L) | Serum effect (mg/L) | FIC index |
| --- | --- | --- | --- | --- | --- | --- | --- | --- | --- | --- |
|  |  | MEM alone | TACN alone | {MEM-TACN4*} | {MEM-TACN8*} | {MEM-TACN16*} | MEM-TACN8* | MEM-TACN8* | MEM-TACN8*/Serum |  |
| Metallo-beta-lactamases |  |  |  |  |  |  |  |  |  |  |
| *E. coli* | NDM-1 | 128 | 1024 | {0.125-4} | {0.125-8} | {0.06-16} | 0.5 | 2 | 0.25 | 0.008 |
| *E. cloacae* | NDM-1 | 16 | >1024 | {{0.06-4} | {0.03-8} | - | 0.125 | 2 | 0.125 | ≤0.005 |
| *C. freundii* | NDM-1 | 8 | 1024 | {0.25-4} | {0.125-8} | {0.125-16} | 0.125 | 1 | 0.25 | 0.023 |
| *P. rettgeri* | NDM-1 | 64 | >1024 | {8-4} | {2-8} | - | 2 | 1 | 2 | ≤0.035 |
| *P. stuartii* | NDM-1 | 16 | >1024 | {2-4} | {2-8} | - | 2 | 1 | 4 | ≤0.253 |
| *E. coli* | NDM-4 | 128 | 1024 | - | {0.125-8} | {0.06-16} | 0.125 | 1 | 1 | 0.008 |
| *E. coli* | VIM-1 | 32 | 1024 | {0.25-4} | {0.125-8} | {0.06-16} | 0.125 | 1 | 0.25 | 0.011 |
| *K. pneumoniae* | VIM-1 | 32 | 512 | {2-4} | {0.125-8} | {0.06-16} | 0.5 | 4 | 8 | 0.019 |
| *E. cloacae* | VIM-1 | 8 | 128 | {1-4} | {1-8} | - | 1 | 1 | 2 | 0.187 |
| *E. coli* | VIM-2 | 8 | 128 | {0.06-4} | {0.03-8} | - | 0.06 | 1 | 0.125 | 0.066 |
| *E. cloacae* | IMP-1 | 32 | 512 | {0.25-4} | {0.125-8} | - | 0.5 | 2 | 2 | 0.019 |
| *K. pneumoniae* | IMP-1 | 32 | 512 | {8-4} | {4-8} | - | 4 | 1 | 16 | 0.141 |
| *E. coli* | IMP-1 | 16 | 512 | {1-4} | {0.06-8} | {0.03-16} | 0.125 | 2 | 0.25 | 0.019 |
| *E. cloacae* | IMP-8 | 8 | 512 | {0.5-4} | {0.125-8} | {0.03-16} | 0.25 | 2 | 0.125 | 0.031 |
| *K. pneumoniae* | IMP-8 | 16 | 1024 | {8-4} | {4-8} | {2-16} | 8 | 2 | 4 | 0.250 |
| Serine-beta-lactamases |  |  |  |  |  |  |  |  |  |  |
| *K. pneumoniae* | OXA-48 | 8 | 128 | {8-8} | {8-16} | {4-64} | NA** | NA | NA | NA |
| *E. cloaeae* | KPC-2 | 32 | 256 |  | {32-32} | {32-64} | NA | NA | NA | NA |
| ATCC susceptible strains |  |  |  |  |  |  |  |  |  |  |
| *K. pneumoniae* ATCC BAA 1706 | Susceptible strain | 0.5 | 1024 | NE*** |  |  | NA | NA | NA | NA |
| *E. coli* ATCC 25922 | Susceptible strain | 0.03 | 1024 | NE |  |  | NA | NA | NA | NA |

\*Fixed concentration of TACN (4, 8, 16 mg/L etc...) with varying concentrations of Meropenem \*\*Not applicable \*\*\* No inhibitory effect

**Table 2:**
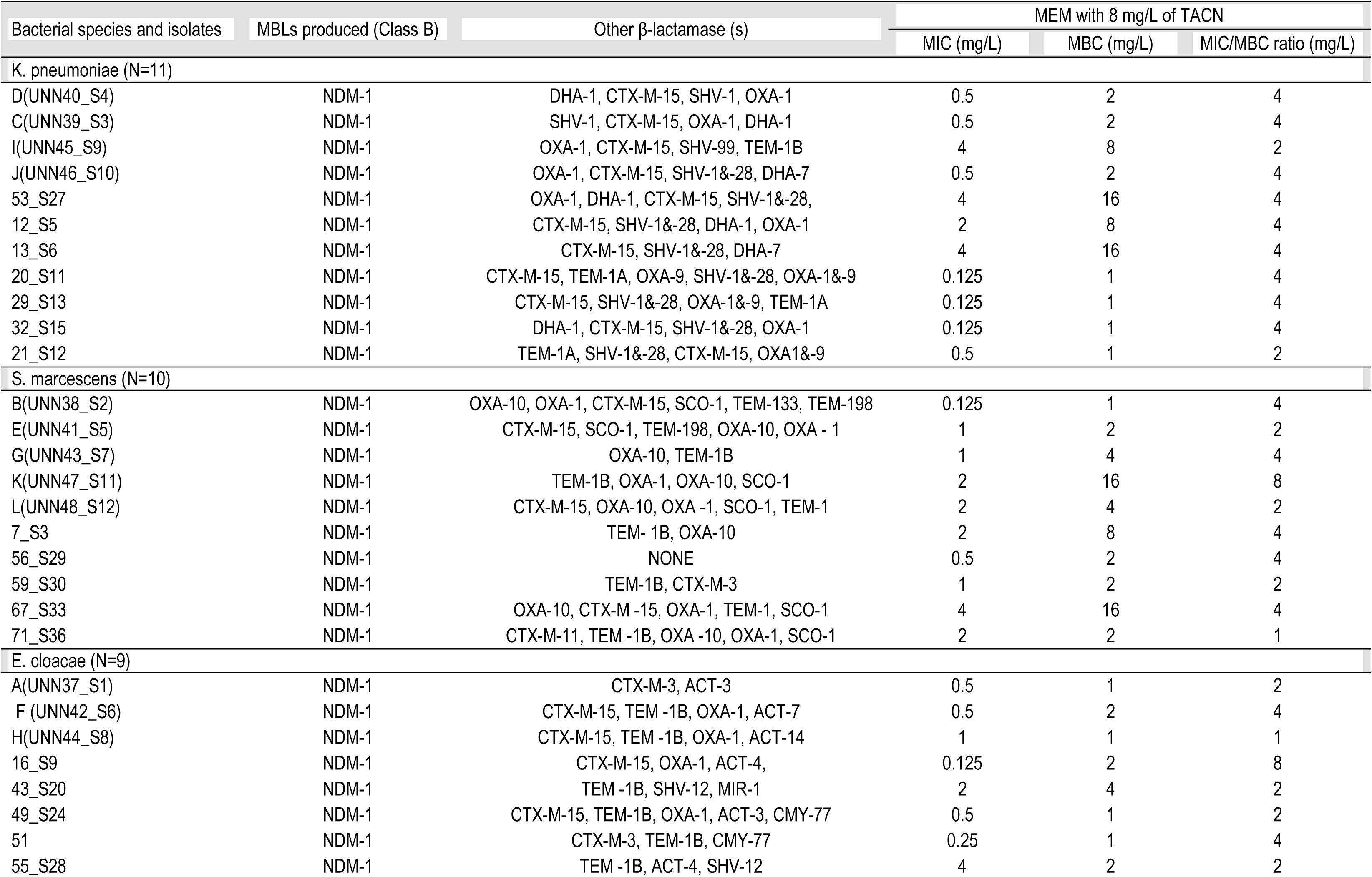
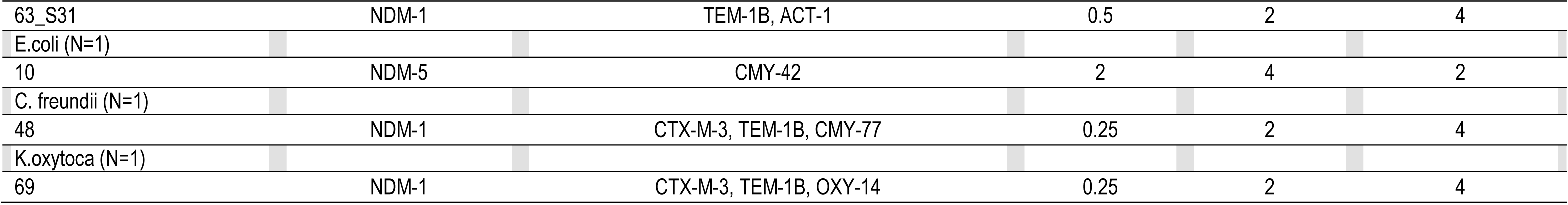
Minimal inhibitory concentrations (MIC) and minimal bactericidal concentrations (MBC) of TACN against South African clinical isolates.

| Bacterial species and isolates | MBLs produced (Class B) | Other β-lactamase (s) | MEM with 8 mg/L of TACN | | |
| --- | --- | --- | --- | --- | --- |
|  |  |  | MIC (mg/L) | MBC (mg/L) | MIC/MBC ratio (mg/L) |
| K. pneumoniae (N=11) |  |  |  |  |  |
| D(UNN40_S4) | NDM-1 | DHA-1, CTX-M-15, SHV-1, OXA-1 | 0.5 | 2 | 4 |
| C(UNN39_S3) | NDM-1 | SHV-1, CTX-M-15, OXA-1, DHA-1 | 0.5 | 2 | 4 |
| I(UNN45_S9) | NDM-1 | OXA-1, CTX-M-15, SHV-99, TEM-1B | 4 | 8 | 2 |
| J(UNN46_S10) | NDM-1 | OXA-1, CTX-M-15, SHV-1&-28, DHA-7 | 0.5 | 2 | 4 |
| 53_S27 | NDM-1 | OXA-1, DHA-1, CTX-M-15, SHV-1&-28, | 4 | 16 | 4 |
| 12_S5 | NDM-1 | CTX-M-15, SHV-1&-28, DHA-1, OXA-1 | 2 | 8 | 4 |
| 13_S6 | NDM-1 | CTX-M-15, SHV-1&-28, DHA-7 | 4 | 16 | 4 |
| 20_S11 | NDM-1 | CTX-M-15, TEM-1A, OXA-9, SHV-1&-28, OXA-1&-9 | 0.125 | 1 | 4 |
| 29_S13 | NDM-1 | CTX-M-15, SHV-1&-28, OXA-1&-9, TEM-1A | 0.125 | 1 | 4 |
| 32_S15 | NDM-1 | DHA-1, CTX-M-15, SHV-1&-28, OXA-1 | 0.125 | 1 | 4 |
| 21_S12 | NDM-1 | TEM-1A, SHV-1&-28, CTX-M-15, OXA1&-9 | 0.5 | 1 | 2 |
| S. marcescens (N=10) |  |  |  |  |  |
| B(UNN38_S2) | NDM-1 | OXA-10, OXA-1, CTX-M-15, SCO-1, TEM-133, TEM-198 | 0.125 | 1 | 4 |
| E(UNN41_S5) | NDM-1 | CTX-M-15, SCO-1, TEM-198, OXA-10, OXA - 1 | 1 | 2 | 2 |
| G(UNN43_S7) | NDM-1 | OXA-10, TEM-1B | 1 | 4 | 4 |
| K(UNN47_S11) | NDM-1 | TEM-1B, OXA-1, OXA-10, SCO-1 | 2 | 16 | 8 |
| L(UNN48_S12) | NDM-1 | CTX-M-15, OXA-10, OXA -1, SCO-1, TEM-1 | 2 | 4 | 2 |
| 7_S3 | NDM-1 | TEM- 1B, OXA-10 | 2 | 8 | 4 |
| 56_S29 | NDM-1 | NONE | 0.5 | 2 | 4 |
| 59_S30 | NDM-1 | TEM-1B, CTX-M-3 | 1 | 2 | 2 |
| 67_S33 | NDM-1 | OXA-10, CTX-M -15, OXA-1, TEM-1, SCO-1 | 4 | 16 | 4 |
| 71_S36 | NDM-1 | CTX-M-11, TEM -1B, OXA -10, OXA-1, SCO-1 | 2 | 2 | 1 |
| E. cloacae (N=9) |  |  |  |  |  |
| A(UNN37_S1) | NDM-1 | CTX-M-3, ACT-3 | 0.5 | 1 | 2 |
| F (UNN42_S6) | NDM-1 | CTX-M-15, TEM -1B, OXA-1, ACT-7 | 0.5 | 2 | 4 |
| H(UNN44_S8) | NDM-1 | CTX-M-15, TEM -1B, OXA-1, ACT-14 | 1 | 1 | 1 |
| 16_S9 | NDM-1 | CTX-M-15, OXA-1, ACT-4, | 0.125 | 2 | 8 |
| 43_S20 | NDM-1 | TEM -1B, SHV-12, MIR-1 | 2 | 4 | 2 |
| 49_S24 | NDM-1 | CTX-M-15, TEM-1B, OXA-1, ACT-3, CMY-77 | 0.5 | 1 | 2 |
| 51 | NDM-1 | CTX-M-3, TEM-1B, CMY-77 | 0.25 | 1 | 4 |
| 55_S28 | NDM-1 | TEM -1B, ACT-4, SHV-12 | 4 | 2 | 2 |
|  |  |  | MIC (mg/L) | MBC (mg/L) | MIC/MBC ratio (mg/L) |
| 63_S31 | NDM-1 | TEM-1B, ACT-1 | 0.5 | 2 | 4 |
| E.coli (N=1) |  |  |  |  |  |
| 10 | NDM-5 | CMY-42 | 2 | 4 | 2 |
| C. freundii (N=1) |  |  |  |  |  |
| 48 | NDM-1 | CTX-M-3, TEM-1B, CMY-77 | 0.25 | 2 | 4 |
| K.oxytoca (N=1) |  |  |  |  |  |
| 69 | NDM-1 | CTX-M-3, TEM-1B, OXY-14 | 0.25 | 2 | 4 |

### Serum effect on MIC

The MIC values of MEM-TACN combinations in the presence of 50% human serum varied from ±1-to 4-fold dilutions compared to that of MEM-TACN in the absence of serum indicating that the inhibition properties of the combination were not affected by the presence of human serum (Table 1).

### Synergistic effect of TACN and MEM on CRE reference strains

The synergy between TACN and MEM against CRE isolates was investigated. In most of the cases, the minimum concentrations of TACN alone required to inhibit the MBL-producing *Enterobacteriaceae* were ≥128 mg/L (Table 1). MEM alone exhibited inhibitory concentrations of ≥8 mg/L, indicating carbapenem-resistant phenotype of these pathogens(17). Challenging these CREs with a combined regimen of MEM-TACN potentiated the effectiveness of MEM. The calculated FIC index of MEM-TACN was ≤0.5, indicating a synergistic effect (Table 1).

### Time-kill kinetics

The combination of MEM-TACN caused a considerable decrease in the CFU/mL relative to the initial bacterial density of approximately 10^6^ CFU/mL) over the increasing time points (0, 1, 2, 4, 6, 8 and 24 h) against the tested isolates at different concentrations of the MIC values evaluated in this study (1x, 4x, 8x MICs) (Figure 2). A 3log_10_ decrease in the CFU/ml was observed when cells were treated with MEM-TACN at 1x, 4x and 8x MICs at 4 h. The time-kill kinetics results indicated that the TACN-MEM combinations had bactericidal activity against the selected CRE isolates. MEM alone when challenged at concentrations of 4x and 8x the MICs no significant decrease was observed during the time interval of 24 h.

**Figure 2:** Time kill kinetics at varying concentrations of MEM and fixed concentration of TACN (8 mg/mL). A) *E. coli* producing NDM-1; B). *E. coli* producing VIM-1; C) *E. cloacae* producing IMP-1 were challenged at 1x, 4x and 8x MIC levels with MEM-TACN combination. The surviving CFU were plated at different time intervals (1, 2, 4, 6, 8, 24 hours). All data points represent the average results for 3 independent experiments.

### Cytotoxicity

The MTT assay was used to measure TACN cytotoxicity (0 –1024 mg/L) in HepG2 cells after 24-hour exposure. A decrease in cell viability was associated with an increase in TACN concentration (Figure 3), and the regression analysis yielded an IC_50_ of 56 mg/L. However, this IC_50_ of TACN was far higher than the fixed MIC value of 8 mg/L used in this study.

**Figure 3:** TACN induced a dose-dependent decrease in HepG2 cell viability following treatment for 24-hour treatment. Higher TACN concentrations resulted in increased cell death.

### Enzymatic assays

TACN efficiently inhibited the activity of NDM-1 in a concentration- and time-dependent manner illustrated by Figure 4A where the effect of increasing concentrations of TACN (0 – 50 µM) on the time-dependent decrease in NDM-1 activity. The kinetic parameters were constructed by nonlinear regression of the K_obs_ versus TACN concentration plot shown in figure 4B. TACN was found to be a potent time-dependent inhibitor with an inhibition constant K_i_ and inactivation constant K_inact_ of 0.044±018 µM and 0.0406 ±0.007 (min^-1^) respectively.

**Figure 4:** A) Time- and concentration-dependent inhibition of NDM-1 by TACN, B) The logarithmic plot of kobs of TACN against TACN concentrations.

### Computational simulations

The fluctuations of enzyme-ligand complex and ligand-associated movements were analyzed to ascertain their structural stability and movements during the simulation. These movements, which ensure their functional reliability and working efficiency, are essential for functionality and existence inside the living system. The overall systems showed appreciable stability as depicted in Figure 6. The root-mean-square deviation (RMSD) of the entire system complex remained stable during the course of simulation of 100 ns.

**Figure 6:** Time-dependent RMSD plots of A) NDM1_TACN,_ B) VIM-2_TACN._ The complexes are represented in black and the free enzymes in red.

The calculated binding free energy of TACN to NDM-1 and VIM-2 by the Molecular Mechanics/Poisson-Boltzmann Surface Area *(MM/GBSA) protocol is shown in Table 3. The calculated contributions favoring the inhibitors’ binding include the electrostatic interactions (Δ Eele), which were −9.4229 kcal/mol for NDM-1_TACN_ complex. The intermolecular van der Waals energy (Δ Evdw) for NDM-1_TACN_ was −46.0898 kcal/mol, also favoring binding. The total binding free energies (Δ Gbind) for NDM-1_TACN_ was −39.1603 kcal/mol. A similar trend was also observed in the complex VIM-2_TACN_ as shown in table 3. The results suggest that the theoretical calculations of the binding free energies support the experimental findings in term of the potentiality of TACN to inhibit the MBL enzymes.

**Table 3:** MM/GBSA binding free energyprofile for NMD1_TACN_, and VIM-2_TCN_

| Binding free energy profile |  |  |  |  |  |
| --- | --- | --- | --- | --- | --- |
| Systems | Evdw | Eelec | $\Delta G_{solv}$ | $\Delta G_{gas}$ | $\Delta G_{bind}$ |
| NDM-1<sub>TACN</sub> complex | -46.0898 | -9.4229 | 16.3523 | -55.5126 | -39.1603 |
| VIM-2<sub>TACN</sub> complex | -50.9300 | -9.9143 | 19.6230 | -63.4178 | -41.2199 |

$\Delta G_{bind}$ - binding energy; $\Delta E_{ele}$ – electrostatic; $\Delta E_{vdw}$ – van der waals; $\Delta E_{gas}$ – gas-phase energy; $\Delta E_{sol}$ – solvation energy. All binding free energy profile are expressed in Kcal/mol.

To gain further understanding of the active-site residue interactions with the compounds, the simulated complexes were observed with LigPlot protein-ligand interaction software. It was observed that the ligand is attached with the active site residues in all complexes as shown in Figure 7. For TACN, His250, Asp124 residues provided H-bonding capability while His122, Asn220 and Trp93 roof the hydrophobic active-site pocket. These residues are providing the hydrophobic surface area while maintaining H-bonding capability.

**Figure 7:** Binding mechanism, illustrated by the interaction of docked ligand (TACN) at the active side of NDM-1 (A) and VIM-2 (B).

## Discussion

This study evaluated the TACN-MEM combination against clinical and reference CRE isolates and identified TACN as a promising MBL inhibitor. The affinity of metal chelators for binding/sequestering metal ions(18) such as zinc at MBLs’ active sites makes them potential inhibitors that can be developed for treating CRE infections. TACN is a metal chelating agent that exhibited no cytotoxic effect at effective concentrations.

We showed that TACN significantly reduced the MIC of MEM, reinstating the carbapenem’s efficacy against 33 South African clinical isolates and 15 MBL-producing reference strains. MEM MICs decreased by 4- to >512-fold changes in the presence of TACN against all the tested isolates, except for SBL producers and susceptible ATCC strains. The SBL-producing *E. cloacae* KPC-2 and *K. pneumoniae* OXA-48 only exhibited 2 or no fold decrease in MEM MICs, showing that the inhibitory effect of TACN was specific for zinc-based β-lactamases. In our previous study NOTA and DPA exhibited similar activity against MBLs at 4 and 8 mg/L(14, 19). Metal chelators are known for their strong affinity toward metal ions(18). Class B MBLs possess zinc at their active sites, thus enabling the TACN to bind or sequester the zinc ions and inhibit the enzymes’ β-lactam hydrolysis activity.

MEM regained its pharmacological properties when combined with TACN. Most of the tested isolates exhibited a MBC/MIC ratio of ≤ 4, except for two clinical isolates, *S. marcescens and E. cloacae*, where the MBC/MIC ratio was 8 (Table 2). Based on the MBC/MIC ratio values, MEM demonstrated its known bactericidal activity in the presence of TACN, and time-kill kinetic studies confirmed the bactericidal property of MEM-TACN combinations. The MEM-TACN combination was synergistic, with an FIC index ranging from ≤ 0.005 to 0.25 for all the CRE reference strains used in the study. Aspergillomarasmine A, a metal chelating agent, was also found synergistic in combination with MEM toward MBLs(20).

The results also showed that serum had no substantial effect on the MICs of TACN-MEM combinations. The minimal or no effects of serum on the activity of MEM was attributed to MEM’s low protein-binding properties(21). An investigation of the effects of serum on TACN or MEM’s MICs may allow speculation on the probable *in vivo* properties of the molecules at pre-clinical stage. Furthermore, TACN did not affect the pharmacological property of MEM, suggesting that TACN could have low plasma protein-binding properties either with or without MEM.

The MTT assay measures the ability of cells to reduce the MTT salt to a formazan product (22). This reaction can only occur in living cells with healthy mitochondria, as it is reliant on NADPH-dependent oxidoreductase enzymes. The cell viability obtained, therefore, reflects the number of viable cells present. TACN exposure resulted in a dose-dependent decrease in cellular metabolic activity and viability in HepG2 cells (Figure 3). However, the IC_50_ obtained was ≥4-fold higher than the TACN inhibitory dose (8 mg/L), and the cell viability was 91% at the upper limit of the inhibitory dose. This cytotoxicity analysis on TACN found the compound possessing tolerable toxicity on HepG2 cell line at the applied concentration, hence showing potential for development as MBLI for clinical use.

An experimental assay together with computational simulation demonstrated an excellent binding affinity between ligand and enzyme complexes as well as potential inhibitory properties. It is known that Class B β-lactamases are susceptible to inhibition by chelating agents during interactions between the chelators (ligands) and the metal ions after ions are released from the proteins (MBL) active site or while still associated with the enzyme complex(23). TACN possessed an apparent potential inhibition property toward NDM-1 enzyme like Aspergillomarasmine A(20) as shown in figure 5. Overall, the calculated variation of RMSD did not show considerable structural shifts, suggesting the stability of the enzyme structure and strength of ligand attachment inside the active site pocket. The binding free energy analysis showed that the intermolecular van der Waals and the electrostatic interaction were the forces driving both systems. However, the total solvation energy (ΔGsol) was unfavorable for all the complexes. Even though hydrogen bonds between the ligand and receptors were observed using LigPlot and chimera (Figure 7), their influences could not compensate for the substantial desolvation penalties during the interaction, thereby leading to the total solvation energy (ΔGsol). Similar results have been reported previously where inhibition was dominated by the van der Waals interactions (24, 25).

**Figure 5:** Residual activity after rapid dilution of enzyme-inhibitor complex. The absorbance changes from the co-incubation of NDM-1 with no inhibitor (•) and NDM-1 + TACN (•) are shown. Slope of the initial rate represents the hydrolytic activity of NDM-1.

In conclusion, this study showed that TACN inhibits Class B1 MBL-producing CREs with minimal cytotoxic effect at effective concentrations. TACN successfully potentiated the antimicrobial properties of MEM as should thus be further investigated as an MBLI for clinical development, in combination with carbapenems. Molecular modeling and docking studies demonstrated the mechanistic pathway of TACN to inhibit NDM-1 and VIM-2 sustaining a stable enzyme-ligand complex with the caveat that this is only simulation of a prediction and not likely to represent the actual binding mode. In-depth analysis and optimization of the metal-chelating agent (TACN), employing a multidisciplinary research approach involving advanced computational simulations, biochemical, microbiological, bio-informatics are needed in order to provide more evidence.

## Materials and methods

### Materials

Meropenem, TACN, dimethyl sulfoxide (DMSO), human serum, cation-adjusted Mueller-Hinton broth (CAMHB) and Mueller-Hinton agar (MHA), phosphate buffered saline (PBS), β-lactamase inhibitor screening kit (MAK222) and 3-(4,5-di methyl thiazol-2-yl)-2,5-diphenyltetrazolium bromide (MTT) were purchased from Sigma Aldrich (St. Louis, MO, USA). Purified NDM-1 was obtained from RayBiotech, Norcross, GA 30092, USA. Immortalized liver carcinoma (HepG2) cells and cell culture consumables were procured from Highveld Biologicals (Johannesburg, South Africa) and Whitehead Scientific (Johannesburg, South Africa) respectively.

## Bacterial isolates

Seventeen carbapenem-resistant *Enterobacteriaceae* reference isolates (comprising 15 metallo-beta-lactamases and two serine-beta-lactamases) obtained from France (Institut Pasteur de France) (Table 1)(26) and thirty-three clinical isolates of metallo-beta-lactamases obtained from Lancet laboratories (private sector, South Africa) were employed in this study (Table 2)(27). *Klebsiella pneumoniae* ATCC BAA 1706 and *Escherichia coli* ATCC 25922 purchased from the American Type Culture Collection (ATCC) were susceptible reference strains used as quality control. All bacterial isolates were fully characterized in previous studies for their phenotypic and genotypic resistance profiles(26, 27). The isolates were sub-cultured twice from freezer stocks onto MHA plates and incubated at 37 °C before the experiments. All subsequent liquid subcultures were derived from colonies isolated from the agar plates and were grown in broth medium (CAMHB).

## Methods

### Minimum Inhibitory Concentrations (MICs), Minimum Bactericidal Concentrations (MBCs)

MICs/MBCs determinations were performed according to CLSI guidelines and protocols described by Keepers *et al.* (17, 28). Briefly, two-fold dilutions of MEM and TACN, from 0.015 to 16 mg/L and 1 to 64 mg/L respectively, were made with CAMHB in 96-well microtiter plates using the checkerboard method(13). A 0.5 McFarland-standardized bacterial inoculum was suspended into 200 μl CAMHB in each microtiter well. The plates were then incubated for 18-22 h at 37°C under aerobic conditions. The MIC was determined as the lowest concentration at which there was no visible growth. Afterwards, an aliquot of 100 μL was taken from the MIC assay wells in which no visible growth was observed and inoculated onto MHA plates for MBC determination by incubating at 37 °C for 24 h. The MBC was determined as the lowest concentration of the test compound that resulted in a ≥99.9% decrease of the initial inoculum in the viable bacterial count on the agar plates. Control wells were filled with the amount of solvent(s) used in dissolving drug candidates, CAMHB and bacteria. Experiments were conducted in triplicate.

### Serum effects on the MIC

The effects of serum on the MIC of TACN-MEM combination were performed using the above-described MIC method. In this assay, however, sterile-filtered serum from human male AB plasma was added to the broth (CAMHB) to prepare 50% human serum in the final culture broth. Reference CRE strains carrying different enzymes (NDM-1, NDM-4, VIM-1, IMP-1 and IMP-8) were used to conduct this experiment.

### Synergistic activity

The Synergistic activity assay was used to describe the effect of compounds (TACN and MEM) working together. The synergistic effect between TACN and MEM was determined by the combination assay as described previously with few modifications(29). Briefly, two-fold serial dilutions of MEM were titrated with a fixed concentration (8 mg/L) of TACN. This 8 mg/L TACN was the lowest concentration that did not inhibit the growth of bacteria alone but considerably potentiated the activity of MEM. MEM and TACN were also tested individually to determine their MICs. The fractional inhibitory concentration (FIC) index was calculated as follows: FIC of MEM (MIC of MEM + TACN/MIC of MEM alone) + FIC of TACN (MIC of MEM + TACN/MIC of TACN alone). The effect/activity of a combination with an FIC index of ≤ 0.5 was considered synergistic while an FIC index of 1 was defined as additive and an FIC index of > 4 was characterized as antagonistic.

### Time-kill assays

Time Kill assays were to measure the quantitative kinetic kill model of TACN for the selected organisms and different time points. Time-kill kinetic studies were conducted referring to the previously reported methods, including those described by CLSI (17, 28). *E. coli* producing *bla*_*NDM-1*_, *E. coli* carrying *bla*_*VIM-1*_ as well as *E. cloacae* producing *bla*_*IMP-1*_ were selected for this assay. Briefly, freshly prepared colonies were re-suspended in 10 mL CAMHB and incubated in an orbital shaking incubator (37°C, 180 rpm) for 1 to 2 h. Cultures were then adjusted to a 0.5 McFarland standard (approximately 1.5x 10^8^ CFU/mL) and further diluted 1:20 in CAMHB so that the starting inoculum was approximately 1.5x 10^6^ CFU/mL. MEM was added to the prepared bacterial suspensions at final concentrations corresponding to 1x, 4x or 8x the MIC of MEM while in combination with TACN at a fixed concentration of 8 mg/L. Also, the time-kill kinetic of the MEM 4x and 8x MIC without TACN was investigated. A growth control with no antibiotic was included in the assay. The initial bacterial load was determined from the growth control test tube instantly after dilution step and was recorded as the CFU count at time zero. The test tubes were incubated in an orbital shaking incubator at 37°C and 180 rpm, and bacterial cell viability counts were performed at 1, 2, 4, 6, 8 and 24 hours by removing 100 μL of the culture, diluting as appropriate, and plating on MHA. The agar plates were incubated at 37°C for at least 22 h. Colonies were counted, and the results were recorded as the number of CFU/mL. A ≥3-log_10_ decrease within 24 hours in the number of CFU/mL was considered bactericidal. The assays were executed in triplicate.

### Cytotoxicity Assay

HepG2 cells were maintained at 37°C in 10% complete culture medium (CCM; Eagles minimum essential medium supplemented with fetal bovine serum, antibiotics and L-glutamine) until confluent. Following trypsinization, 1.5x 10^4^cells/well were allowed to adhere to a 96-well plate overnight. The cells were treated with seven dilutions of TACN (16 – 1024 mg/L) prepared in CCM. Untreated cells (CCM only) served as the control. After 24 hours the treatment was replaced with 20µl MTT solution (5mg/mL MTT in PBS) and 100µl CCM for 4 hours. The resulting formazan product was solubilized in 100µl DMSO (1 hour), and the absorbance at 570nm/690nm was determined (BioTek µQuant plate reader; BioTek Instruments Inc., USA). The average absorbance values were used to calculate cell viability 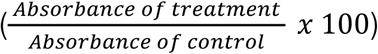 and determine the IC_50_ (GraphPad Prism^®^ v5.01). All experiments were performed in triplicate.

### Enzymatic assays: Enzyme inhibition and inactivation kinetic parameters

The inhibitory effect of TACN was monitored by observing the rate of hydrolysis of the chromogenic nitrocefin, using purified preparations of sub-class B1 MBL enzyme (NDM-1) obtained from Raybiotech (Norcross, GA 30092, USA). The assays were performed using the β-lactamase inhibitor screening kit’s (MAK222) according to the manufacturer’s instructions. Briefly, 5 nM of NDM-1 was pre-incubated with various concentrations of TACN varying from 0 - 50 µM at 25°C in a final experimental volume of 100 μL of β-lactamase assay buffer before adding nitrocefin. The nitrocefin hydrolysis was monitored at 25°C by following the absorbance variation at 490nm using a plate reader spectrophotometer (SPECTROstar^Nano^ BMG Labtech, Germany). The residual activity was determined and compared to that from the pre-incubation of the enzyme with the same procedure but without TACN. The effect of TACN-mediated inactivation of NDM-1 was determined by plotting the natural logarithm (LN)-linear plot of the percentage of the remaining enzyme residual activity versus the pre-incubation time, and the observed initial inactivation rate constant (K_obs_) was calculated from the pseudo-first-order kinetic slope. The rate constant for enzyme inactivation (k_inact_) and the inactivation constant (K_I_) were calculated (GraphPad Prism5; Software Inc., San Diego, CA) according to the following hyperbolic equation as previously described by Wong *et al.* (30):

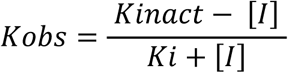

Where *K*_*obs*_ is the initial rate constant for TACN inactivation and [I] is the initial inhibitor concentration. The preliminary mode of inhibition of NDM-1 by TACN was also assessed using the β-lactamase inhibitor screening kit reagents. Briefly, 500 nM of purified NDM-1 was pre-incubated with 20 μM of TACN at 25°C for 20 min, followed by rapid dilution with the addition of 30 μM of nitrocefin; after which, changes in absorbance were monitored at 490 nm for 30 min. Experiments were conducted in triplicate.

### Molecular modelling

#### System preparation

The crystal structures (PDB ID: 3Q6X, and 5ACU) were retrieved from the RSCB Protein Data Bank (https://www.rcsb.org/pdb/). The missing residues were added using a graphical user interface of Chimera, a molecular modelling tool(31). A ligand interaction map was generated using the web version of PoseView(32).

#### Molecular Docking

Docking calculations were obtained using AutoDock Vina software(33). Geister partial charges were assigned, and the AutoDock atom types were defined using the AutoDock graphical user interface supplied by MGL tools(34). The docked conformations were generated using the Lamarckian genetic algorithm (LGA)(35). The binding affinities calculated for each of the two structures (enzyme: inhibitor complexes) were expressed in kcal/mol. This technique has been validated in previous studies(36). The grid box was defined using Autodock Vina with the grid parameters being X= 26, Y= 18 and Z= 20 for the dimensions and X= −2.502, Y= −5.255 and Z= 32.723 for the center grid box. After using Autodock Vina, the ten conformations with lowest binding energy were chosen for molecular docking.

#### Molecular dynamics simulation

Missing parameters for the ligand in the Cornell et al. force field were created (37) in the absence of available parameters. Optimization of the ligands was first performed at the HF/6-31G* level with the Gaussian 03 package. The restrained electrostatic potential (RESP) procedure(38) was used to calculate the partial atomic charges. General amber force GAFF(39) force field parameters and RESP partial charges were assigned using the ANTECHAMBER module in the Amber14 package. Hydrogen atoms of the proteins were added using the Leap module in Amber12. The standard AMBER force field for bioorganic systems (ff03) was used to define the enzyme parameters. Counter ions were added to neutralize the enzyme charge. The system was enveloped in a box of equilibrated TIP3P water molecules with 8 Å distance around the enzyme. Cubic periodic boundary conditions were imposed, and the long-range electrostatic interactions were treated with the particle-mesh Ewald method (40) implemented in Amber12 with a non-bonding cut-off distance of 10 Å.

Initial energy minimization, with a restraint potential of 2 kcal/mol Å^2^ applied to the solute, was carried out using the steepest descent method in Amber12 for 1000 iterations followed by conjugate gradient protocol for 2000 steps. The entire system was then freely minimized for 1000 iterations. Harmonic restraints with force constants 5 kcal/mol Å^2^ were applied to all solute atoms during the heating phase. A canonical ensemble constant number (N), volume (V), and temperature (NVT) MD was carried out for 50 ps, during which the system was gradually annealed from 0 to 300 K using a Langevin thermostat with a coupling coefficient of 1/ps. Subsequently, the system was equilibrated at 300 K with a 2 fs time step for 100 ps while maintaining the force constants on the restrained solute. The SHAKE algorithm(41) was employed on all atoms covalently bonded to a hydrogen atom during equilibration and production runs. A production run was performed for 2 ns in an isothermal-isobaric (NPT) ensemble using a Berendsen barostat(42) with a target pressure of 1 bar and a pressure coupling constant of 2 ps with no restraints imposed. The coordinate file was saved every 1 ps, and the trajectory was analyzed every 1 ps using Ptraj module implemented in Amber14.

## Author contributions

Co-conceptualized the study: AMS, DGA, JOS, LAB and SYE. Performed the experiments: AMS, DGA, HMK and RK. Analyzed the data: AMS, DGA, HMK and RK. Vetting of the results: All. Wrote the paper: AMS. Undertook critical revision of the manuscript: All.

### Acknowledgement

We are grateful to the South African National Research Foundation (**Grant No.: 85595** awarded to Professor S.Y. Essack as Incentive Funding for Rated Researchers) and College of Health Sciences, UKZN, for funding this study.

